# Single-nucleotide resolution mapping of m^6^A of zebrafish mRNAs in early embryonic development links m^6^A modifications to the maternal-to-zygotic transition

**DOI:** 10.1101/2025.11.19.688951

**Authors:** Alex Wardale, Yong Peng, Ruiqi Ge, Tor Erik Jørgensen, Steinar Daae Johansen, Chuan He, Igor Babiak

## Abstract

N^6^-methyladenosine (m^6^A) is the most abundant epigenetic modification on eukaryotic mRNA and has important roles in gene expression. While m^6^A distribution and localization have been defined in zebrafish using antibody-based methods, no research has been published on the distribution of m^6^A during zebrafish embryogenesis with single-base precision. Here, we present the transcriptome-wide m^6^A landscape at single-nucleotide resolution during zebrafish early embryonic development using m^6^A-selective allyl chemical labeling and sequencing (m^6^A-SAC-seq). We identify over 36,000 high-confidence unique m^6^A sites that are primarily located in the DRACH consensus motif in long internal exons, 3’UTRs, and near stop codons. Our results show a >4-fold decrease in m^6^A levels from 256-cell to 50% epiboly, clearly linking this methylation to zebrafish maternal-to-zygotic transition. We highlight that differential methylation between developmental stages was observed in transcripts associated with mRNA metabolism, translational processing, and organ development.

## Introduction

Over 170 different post-transcriptional RNA modifications have been identified (Cappannini et al. 2024), with several of them being involved in regulation of gene expression (Arzumanian et al. 2022). However, the discovery of FTO alpha-ketoglutarate dependent dioxygenase (FTO) and alkB homolog 5 (ALKBH5) as “erasers” of N^6^-methyladenosine (m^6^A) led to the exponential increase in research dedicated to this reversible modification (Jia et al. 2011; Zheng et al. 2013). Since then, m^6^A has been identified and described in a wide variety of organisms including yeast, plants, insects and mammals (Bodi et al. 2010; Dominissini et al. 2012; Lence et al. 2016; Wang et al. 2024) and has been defined as the most abundant posttranscriptional modification on eukaryotic mRNA (Dominissini et al. 2012). Furthermore, m^6^A is important in gene expression processes, marking transcripts for translation and degradation, among other roles (Wang et al. 2015).

By implementing modified Next Generation Sequencing (NGS) library preparation protocols, researchers have been able to describe m^6^A stoichiometry and distribution across transcriptomes, identifying the DRACH (D = G, A or U; R = A or G; H = A, C or U) consensus motif, and a preference towards 3’UTRs, near stop codons and in long internal exons (Dominissini et al. 2012). Initially, these protocols presented several drawbacks, namely low resolution in a 100-200 nucleotide window, or high input material requirements upward of 5 µg of poly(A)-enriched mRNA (Linder et al. 2015; Meyer et al. 2012; Zhang et al. 2019; Wang et al. 2020). To address these limitations, an enzyme conversion method was developed that required as little as 2 ng of poly(A)-mRNA, known as m^6^A-selective allyl chemical labeling and sequencing (m^6^A-SAC-seq; Ge et al. 2022; Hu et al. 2022). This stringent method labels and converts m^6^A, which, in combination with its low input material requirement, allows for single-nucleotide detection accuracy of this methylation transcriptome-wide.

In zebrafish, m^6^A had been initially linked to the clearance of maternal mRNAs (Zhao et al. 2017); however, whether this process is reliant on a specific subset of m^6^A “readers” has been questioned and contrasted (Kontur et al. 2020). Additional unpublished research carried out on zebrafish early embryogenesis revealed a relation between m^6^A methylation and the developmental process of zygotic genome activation, ZGA (Aanes et al. 2019). Most recently, single-cell m^6^A mapping was carried out on zebrafish embryos, detecting an enrichment for the modification near stop codons and at its predominant motifs (Li et al. 2024). Nevertheless, the aforementioned investigations have struggled to both precisely describe m^6^A location and its dynamics during early zebrafish development.

To address this, we set out to map the m^6^A methylome at single-base resolution during zebrafish early embryonic development. By implementing m^6^A-SAC-seq, we sequenced embryo transcriptomes from zygote to gastrula period, encompassing the process of MZT, and identified m^6^A sites transcriptome-wide. These results are unprecedented, as m^6^A has not been previously described at single-nucleotide resolution during the embryonic development of zebrafish. Finally, this study contextualizes zebrafish in the conserved transcriptomic distribution of m^6^A methylations alongside other vertebrate early embryonic m^6^A localization.

## Results

HiSeq paired-end sequencing resulted in an average of 50 million reads for each WT and FTO+ replicate (Supplementary Table S2). On average, around 50% of the reads were mapped to the zebrafish GRCz11 genome.

The analysis of downstream data from m^6^A-SAC-seq revealed a total of 36,291 unique m^6^A sites throughout early development, with 147 of them being shared across all the investigated developmental stages (Figure 1A; Supplementary Files S1 & S2). The total number of high-confidence m^6^A sites identified in each stage was: 12,104 in 1-cell, 15,264 in 256-cell, 13,681 in oblong, 3,531 in 50% epiboly and 2,165 in bud (Figure 1A). The proportion of non-DRACH/DRACH sites corresponding to each stage was 0.34, 0.39, 0.29, 0.52 and 0.80, respectively (Figure 1B). On average, 83% of m^6^A sites were located on genic regions, and over half of these mapping to a unique transcript (Supplementary Table S3).

**Figure 1:**
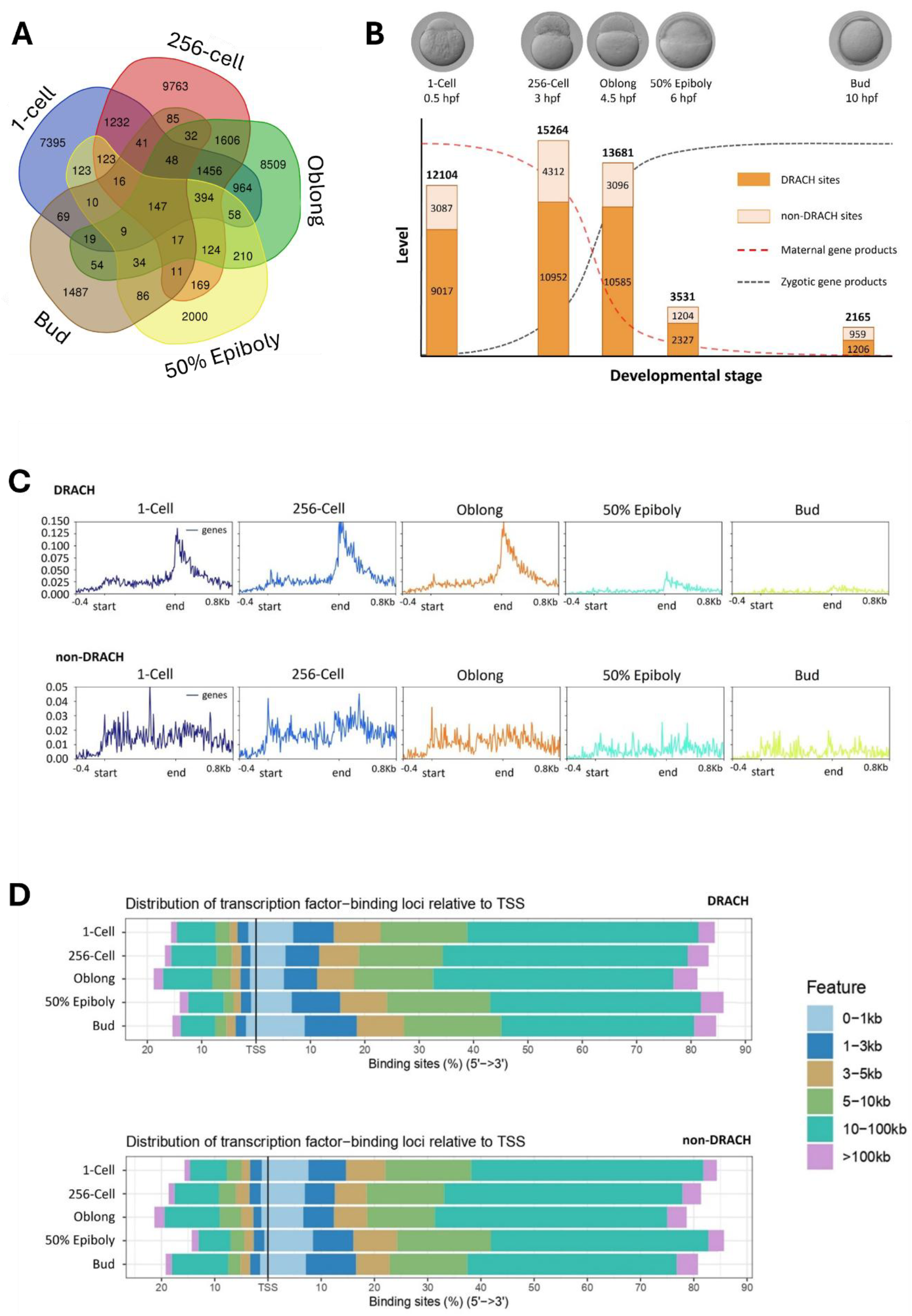
**A**. Number of shared nucleotide-specific m^6^A locations between different zebrafish embryonic development stages. **B**. Representation of the number of m^6^A sites present in different zebrafish embryonic stages in the context of maternal and zygotic gene product levels (the latter adapted from Laue et al. 2019). Bars are divided into number of DRACH (D = G, A or U; R = A or G; H = A, C or U) and non-DRACH m^6^A sites per developmental stage. Embryo photographs by courtesy of Dr. Christopher Presslauer, Nord University. **C**. Overall frequency of m^6^A methylations across genes for DRACH and non-DRACH sites. **D**. Percentage distribution of DRACH and non-DRACH m^6^A sites with relative distance to the transcription start site (TSS) of a gene.

DRACH m^6^A sites were primarily located near stop codons, with a predominance of sites being located 10-100kb downstream of TSS for all embryonic stages (Figure 1C). On the other hand, non-DRACH sites were not predominant near stop codons, although they were also primarily located 10-100kb downstream of the TSS. Within transcripts, m^6^A sites were primarily localized in 3’UTRs and exonic regions throughout development, regardless of the consensus motif (Figure 1D). For DRACH sites, the GGACU motif was the most frequent, while AGAUA was the most common for non-DRACH motifs.

Methylated fraction distribution of m^6^A locations differed for DRACH and non-DRACH motifs (Figure 2A). Sites located in a DRACH motif had higher likelihood of presenting >90% methylated fraction (probability density >0.02), while non-DRACH counterparts were predominantly within 25-60% methylated and >90% methylated fractions (probability density >0.01). These distributions were further investigated, to see if there was any relation between methylated fraction and location of m^6^A sites on mRNA transcripts; however, no link was found between the two (Supplementary Figures S1 - S4).

**Figure 2:**
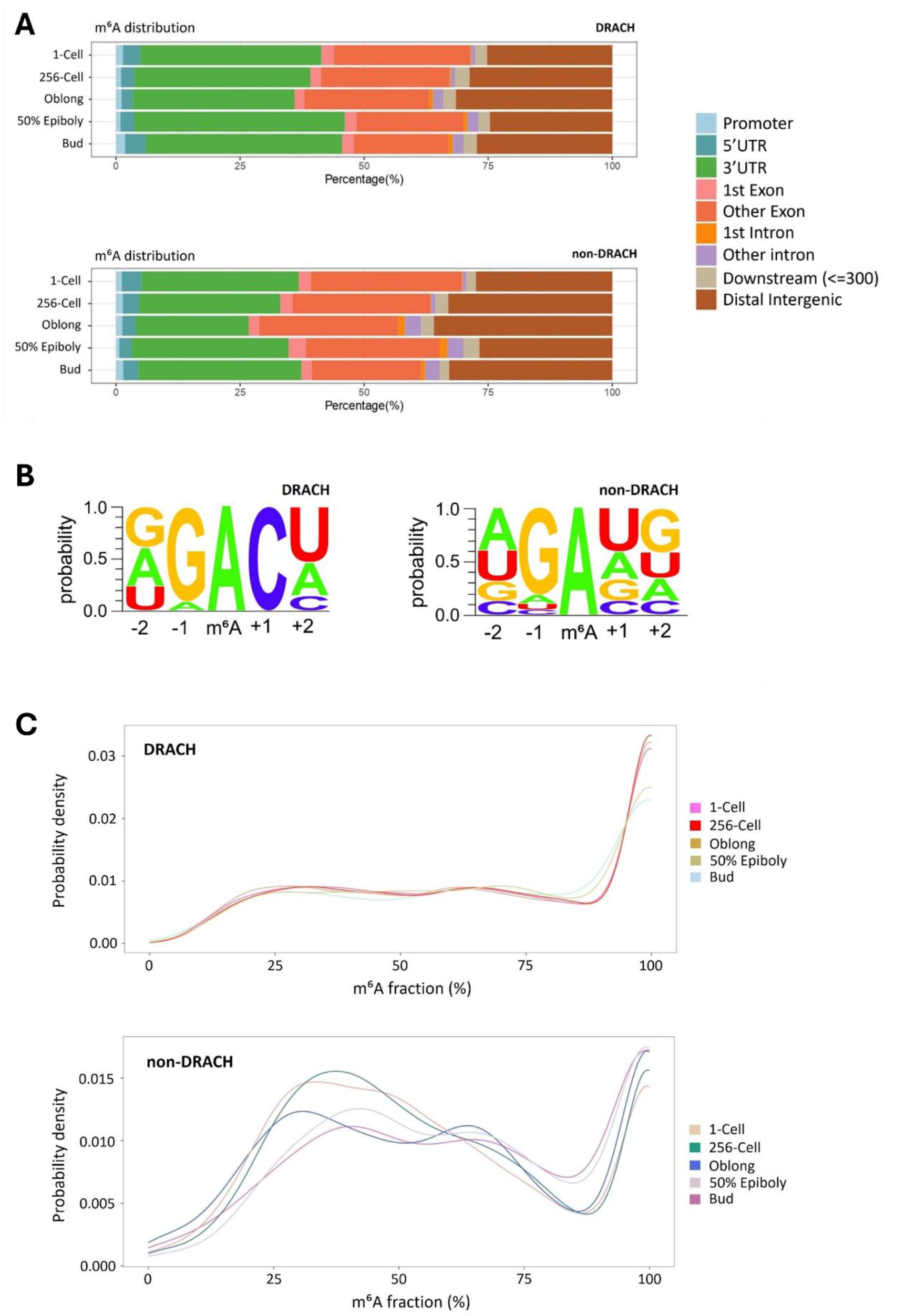
**A**. Percentage distributions of DRACH and non-DRACH m^6^A locations across genomic regions. **B**. Probability of each individual DRACH and non-DRACH motif considering the central A as the m^6^A-modified site. **C**. DRACH and non-DRACH m^6^A methylation stoichiometry.

To avoid confusion of m^6^A sites presenting ambiguous or unambiguous mapping, we investigated the locations relative to the parent gene. Gene ontology (GO) analyses were carried out for differentially methylated genes between developmental stages (Supplementary File S3). Differentially methylated genes were categorized depending on whether they presented an increase (DMF+) or decrease (DMF-) in methylated fractions when comparing between stages. The analysis of DMF-showed an enrichment in, among others, RNA methylation, modification and 3’-end processing categories, alongside mRNA splicing and stability categories throughout development (Supplementary Figure S5; Supplementary File S4). Likewise, increased fold enrichment was observed for methyltransferase complex and nucleus-related categories in regard to cellular localization, while RNA binding, mRNA binding and methyltransferase activity categories were overrepresented in terms of function. On the other hand, GO analysis of genes presenting DMF+ were more commonly related to embryonic development processes, such as mitotic nuclear division and cell cycle phase, somitogenesis and gastrulation, among others (Supplementary Figure S6; Supplementary File S5). When it comes to localization and function, DMF+ presented an enrichment for nucleus and spliceosomal complex-related categories and mRNA binding and ribosomal-related categories, respectively.

In addition to DMF- and DMF+ pairwise comparisons between developmental stages, a GO analysis was carried out on genes that were differentially methylated during MZT by comparing the two earlier embryonic stages to the two later stages (Supplementary File S6). Overrepresented categories for DMF-genes were mainly related to mRNA transport and 3’UTR binding, RNA methylation and binding, and translation initiation categories, while their DMF+ counterparts were enriched for developmental processes such as mesoderm, otic vesicle and digestive tract development, with transcription and translation-related categories enriched in terms of localization and function (Supplementary Figure S7; Supplementary File S7).

A total of 512 genes showed >50% methylated fraction throughout all the investigated developmental stages (Supplementary File S8). They were overrepresented in RNA splicing, modification-dependent processes (Supplementary Figure S8; Supplementary File S9). In addition, the 147 m^6^A sites that were conservatively methylated throughout development (Figure 1A; Supplementary File S2), were analyzed. GO terms enriched included mRNA splicing, alternative mRNA splicing, mRNA binding and embryonic morphogenesis, with a notable >8-fold increase for mRNA categories and >4-fold increase for the embryonic category (Supplementary File S10).

## Discussion

In vertebrates, m^6^A has been primarily linked to mRNA metabolism, namely transcript degradation and increased translational efficiency (Wang et al. 2015). Previous studies on m^6^A methylomes have revealed a high conservation of this modification among eukaryotic mRNAs (Dominissini et al. 2012; Li et al. 2024; Liu et al. 2021; Wang et al. 2024). We demonstrate that the general distribution and localization of m^6^A is conserved between zebrafish and other vertebrates, as we observe a higher abundance of m^6^A sites in long internal exons, 3’UTRs, and near stop codons (Figure 1C & 1D). Furthermore, m^6^A sites are primarily located in the DRACH consensus motif throughout zebrafish development, namely the GGACU motif (Figure 2B), which is in line with research in human and mouse cell lines (Dominissini et al. 2012), zebrafish zygotes (Li et al. 2024), and bovine preimplantation development (Iyyappan et al. 2025).

In animals, MZT is characterized by two main processes: degradation of maternal mRNAs and transcription activation of zygotic mRNAs (Walser & Lipshitz, 2011). In zebrafish, MZT occurs around 3 hours post fertilization (hpf), much later than its mouse and human counterparts, which take place after the first and second cellular division, respectively (Laue et al. 2019; Tesarik 2022). We find a substantial increase in m^6^A sites during the first two hours of development, with a steep decline in m^6^A levels after the oblong stage, and a significant loss of modified sites from 3 – 6 hpf (Figure 1B). Our results clearly link the m^6^A modifications of mRNAs to the process of MZT. These results are supported in a previous unpublished study on zebrafish embryogenesis using an antibody-based approach in mapping m^6^A during zebrafish development at lower positional and stoichiometric resolution (Aanes et al. 2019). Furthermore, our finding of a steep decline in m^6^A levels after oblong corroborates previous observations in zebrafish (Kontur et al. 2020; Zhao et al. 2017). On the other hand, the increase in m^6^A sites during the first two hours of development is intriguing, as the genome is transcriptionally inactive during the initial stages of embryogenesis, with the exception of micro RNA miR-430, and a subset of other ncRNAs and mRNAs (Giraldez et al. 2006; Heyn et al. 2014; Wu & Vastenhouw, 2020). The increase in m^6^A levels from 1-cell to 256-cell stage could be linked to an m^6^A-dependent stabilization of mRNAs, whereby m^6^A-methylated transcripts are selectively bound to by mRNA-binding proteins to enhance transcript stability (Huang et al. 2018). This would suggest that non-m^6^A-methylated mRNAs are potentially degraded faster, allowing the capture of a higher number m^6^A-methylated transcripts, and explaining the increase in m^6^A sites identified in the 256-cell stage.

The first rounds of cellular division after zygote formation in zebrafish involve the initiation of many regulatory processes such as posttranscriptional and translational regulation (Chang et al. 2018; Lee et al. 2013). Post-MZT, we observed a notable decrease in m^6^A levels in genes that are involved in these processes, mainly translational initiation, mRNA transport and RNA binding (Figure 7C). It is possible that the incorporation of m^6^A methylated maternal mRNAs may be crucial to provide a kickstart to these regulatory processes. This hypothesis is supported by the role of this modification in zebrafish mRNA metabolism, as m^6^A can serve as a signal to enhance translation or mark for decay, among others (Kontur et al. 2020). Maternal transcripts harboring increased levels of m^6^A could be preferentially translated while also being submitted to the processes of deadenylation and degradation in parallel, meaning that their processing is faster while also presenting shorter half-lives.

During the gastrulation period of zebrafish embryogenesis (5 - 10 hpf), processes that are essential for correct development such as the completion of epiboly and the production of germ layers and embryonic axis take place (Kimmel et al. 1995). In addition, zebrafish gastrulation is mainly controlled by the zygotic genome and is also dependent on the successful clearance of the maternal gene products (reviewed in Solnica-Krezel 2020). We observe an increase in m^6^A levels in transcripts originating from genes related to developmental processes in 50% epiboly and bud stage, including cell migration involved in gastrulation, and otic vesicle, digestive tract and mesoderm development, among others (Figure 7C). Although tempted to speculate that gastrulation-related RNAs become methylated to signal for relevant processing, this could also be due to the sudden shift in transcript content, as zygotically transcribed mRNAs, which can have a different m^6^A signature, begin to surpass the number of maternal transcripts (Aanes et al. 2011). Furthermore, primary organogenesis is a process that occurs post-bud stage (Kimmel et al. 1995), suggesting that increased methylated fractions in transcripts related to organ development processes could be linked to a preparative role of m^6^A, priming mRNA transcripts for downstream translation.

The conserved tendency of m^6^A distribution across eukaryotes along with the dynamic and reversible nature of this modification has been one of the highlights of methylome studies for over a decade (Dominissini et al. 2012; Jia et al. 2011; Meyer et al. 2012). Our results show that not only is m^6^A dynamic in terms of levels of methylation during early zebrafish development, but also in number of m^6^A sites. It shows that during zebrafish embryogenesis, although non-random in terms of transcript region and sequence motif frequencies, m^6^A locations are highly dynamic in terms of site-specific location and stoichiometry. These results suggest that the role of m^6^A is likely related to the global position on a transcript rather than a specific nucleotide needing to be methylated to carry out its function.

## Acknowledgements

The authors would like to thank C. R. Amundsen and S. E. Butwell for their support in the zebrafish research facility, and L. M. Jakt for his input in bioinformatic discussions. This study was funded by the Research Council of Norway, InnControl project (grant nr. 275786).

## Author Contributions

A.W. performed the investigation and experiments, data visualization, and wrote the manuscript. R.G. carried out library preparation and sequencing. Y.P. lead data processing and visualization. T.E.J. and S.D.J. supervised the project and provided feedback on the manuscript. C.H. conceived the project, provided access to state-of-the-art methods and supervised the investigation. I.B. conceived the project, secured funding, supervised the investigation, and provided feedback on the manuscript.

## Data Availability Statement

The raw sequencing datasets generated in this study have been deposited in the NCBI Sequence Read Archive under BioProject PRJNA1364744 (SRA accessions SRR36031735-SRR36031764). All other data supporting the findings of this study are available within the article and its Supplementary Information files.

