## Supplementary Information (Materials & Methods, Supplementary Figures, Supplementary Data File Legends) for "Single-nucleotide resolution mapping of m^6^A of zebrafish mRNAs in early embryonic development links m^6^A modifications to the maternal-to-zygotic transition"

#### Materials and Methods

##### Zebrafish husbandry and embryo collection

All procedures involving zebrafish were conducted in compliance with the Norwegian Regulation on Animal Experimentation (The Norwegian Animal Protection Act 20 December 1974, no. 73). Housing, husbandry and handling were approved by the Norwegian Animal Research Authority (Utvalg for forsøk med dyr, forsøksdyrutvalget, Norway) General License for Fish Maintenance and Breeding (Godkjenning av avdeling for forsøksdyr) no. 17 and approved by the Nord University (Norway) ethics committee. Fish were kept in a Tecniplast recirculating water system (Scanbur A/S, Karlslunde, Denmark) at 28.5°C with a standard photoperiod regime (14 h light – 10 h dark) at Mørkvedbukta Research Station (Bodø, Norway). For spawning, adults were kept in 1.7 L slope breeding tanks overnight in groups of 2-3 males and 2-3 females per tank. Embryos were collected ~15 min after onset of the light period and were incubated in a Sanyo MIR-254-PE refrigerated incubator (Sanyo Electric, Osaka, Japan). Embryonic developmental stages were identified under a stereo microscope according to (Kimmel et al. 1995). Embryos were sampled at the following stages: 1-cell, 256-cell, oblong, 50% epiboly, and bud. Sampled embryos were collected in batches of 30 embryos in 2 mL cryotubes and snap frozen in liquid nitrogen.

##### RNA isolation and ribodepletion

Total RNA was extracted using QIAzol® Lysis Reagent (Qiagen, Hilden, Germany. Cat. No. 79306) following manufacturer protocol. Ribodepletion was carried out using riboPOOLs (probes) following manufacturer instructions (siTOOLS Biotech, Planegg, Germany) and Dynabeads MyOne Streptavidin C1 magnetic beads (Thermo Fisher, Massachusetts,

United States. Cat. No. 65001). RNA concentration was measured using a 2200 TapeStation System (Agilent Technologies, California, United States) and Qubit™ 4 Fluorometer (Thermo Fisher, cat. No. Q33238) both pre- and post-ribodepletion (Supplementary Table S1).

**Library construction and sequencing**

Three replicates per developmental stage of a minimum of 30 ng ribo-depleted total RNA per sample were purified and fragmented as carried out in Hu et al. 2022. As per m$^6$A-SAC-seq experiments, three separate libraries per sample were prepared: an untreated reference input library, an experimental library (FTO$^-$) with allylic-m$^6$A conversion, and a background noise control library (FTO$^+$) with m$^6$A methylation removal. Three prime (3')-adapter ligation, m$^6$A labeling by MjDim1 and final library preparation, purification and Illumina HiSeq sequencing were conducted as described in Hu et al. 2022.

**Data analysis and m$^6$A site calling**

Preprocessing of m$^6$A-SAC-seq data, mapping and m$^6$A site calling were carried out using the source codes available at https://github.com/CTLife/m6A-SAC-seq and https://github.com/shunliubio/m6A-SAC-seq, using zebrafish as the reference genome (GRCz11 available at https://www.ensembl.org/Danio_rerio/). Stringent m$^6$A site calling was implemented by only keeping DRACH sites with background mutation rates of < 5%, mutation rates of > 5%, *P* value < 0.1 and a minimum coverage of 4. For non-DRACH sites, m$^6$A was only identified if background mutation rates < 5%, mutation rates > 10%, *P* value < 0.001 and a minimum coverage of 11. Final m$^6$A sites were only called if they were identified in at least two of the three replicates and were considered as high-confidence m$^6$A sites (Hu et al. 2022).

**$m^6A$ region distribution and differential methylation analysis**

To calculate differential methylation by stage in pairwise fashion, we used the following formula:

$$\Delta\, methylation\ G_{X\ Stage\ A-Stage\ B} = \left(\frac{\sum G_X\ methylated\ fractions}{max\ nr\ m^6A\ sites\ G_X}\right)_{Stage\ B} - \left(\frac{\sum G_X\ methylated\ fractions}{max\ nr\ m^6A\ sites\ G_X}\right)_{Stage\ A}$$

where *G* stands for gene, *Stage* refers to developmental stage, and *max nr $m^6A$ sites* correspond to the highest number of $m^6A$ locations identified in a specific transcript for any of the five embryonic stages. The cutoff for differentially methylated transcripts was defined as a change of 20%, for both positive (DMF+) and negative (DMF-) differentially methylated fractions.

**Gene Ontology**

Gene ontology was carried out for differentially methylated genes using Panther.db to identify associated biological processes, cellular components and molecular functions using Fisher's Exact statistical overrepresentation test with *Danio rerio* as the reference organism for the whole-genome list, using false discovery rate as the correction method (Mi et al. 2019; Thomas et al. 2022).

**Graphical representations**

Representation of common $m^6A$ sites between developmental stages was carried out using the Venn Diagram tool offered by the Bioinformatics & Evolutionary Genomics group at University of Gent (available at https://bioinformatics.psb.ugent.be/webtools/Venn/).

$m^6A$ site distribution for DRACH and non-DRACH motifs relative to start and end of transcripts were generated using deepTools (Ramírez et al. 2016). Global genomic distribution of $m^6A$ locations and relative to transcription start sites (TSS) were generated by ChIPseeker (Wang et al. 2022; Yu et al. 2015). Methylated fraction density distribution of DRACH and non-DRACH motif $m^6A$ sites, and *k*-means clustering of $m^6A$ sites were plotted using in-house scripts.

For enriched sequence motifs, WebLogo 3 was used to generate *5-mer* plots that encompassed $m^6A$ methylated sites across the transcriptome per stage using the RNA sequence type (available at https://weblogo.threeplusone.com/).

### Supplementary Figures

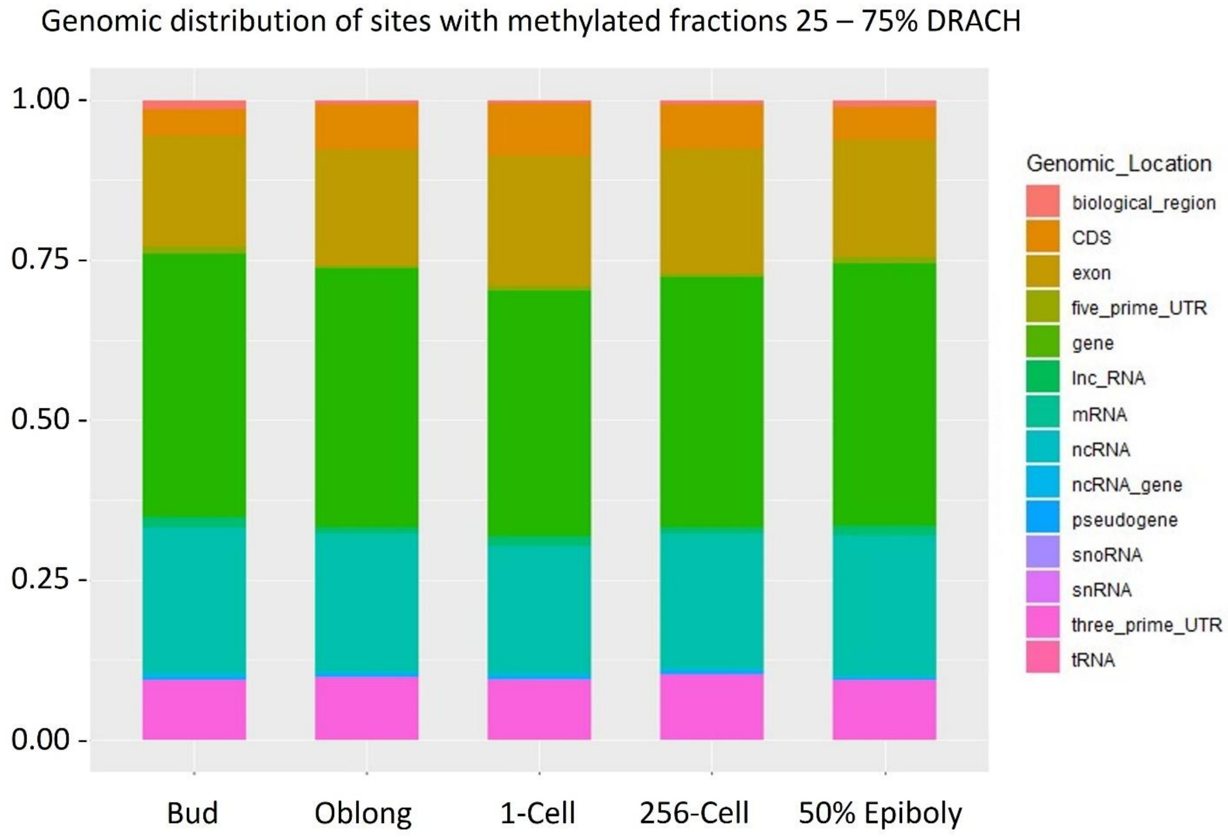


**Supplementary Figure S1:** Genomic distribution of DRACH m$^6$A sites that present 25% - 75% methylated fraction.

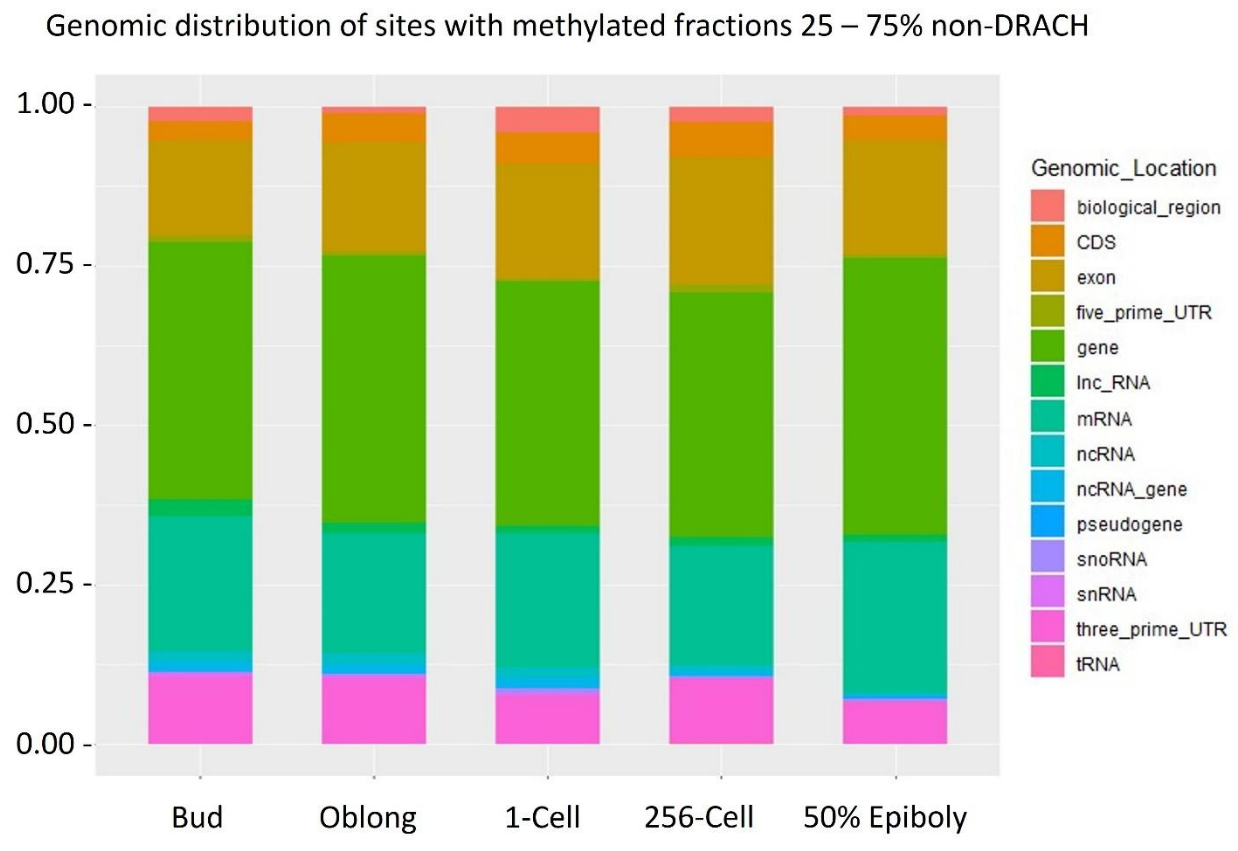


**Supplementary Figure S2:** Genomic distribution of non-DRACH $m^6A$ sites that present 25% - 75% methylated fraction.

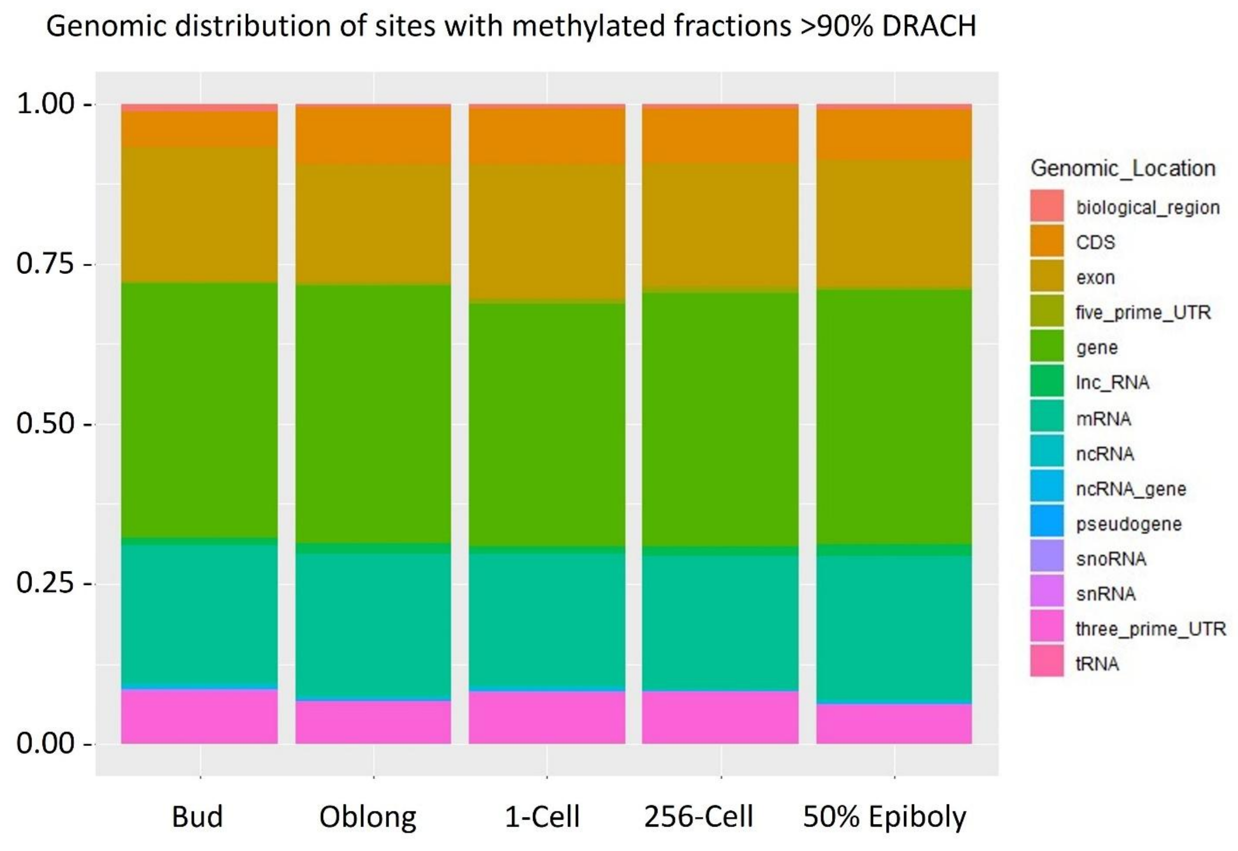


**Supplementary Figure S3:** Genomic distribution of DRACH m$^6$A sites that present 90% - 100% methylated fraction.

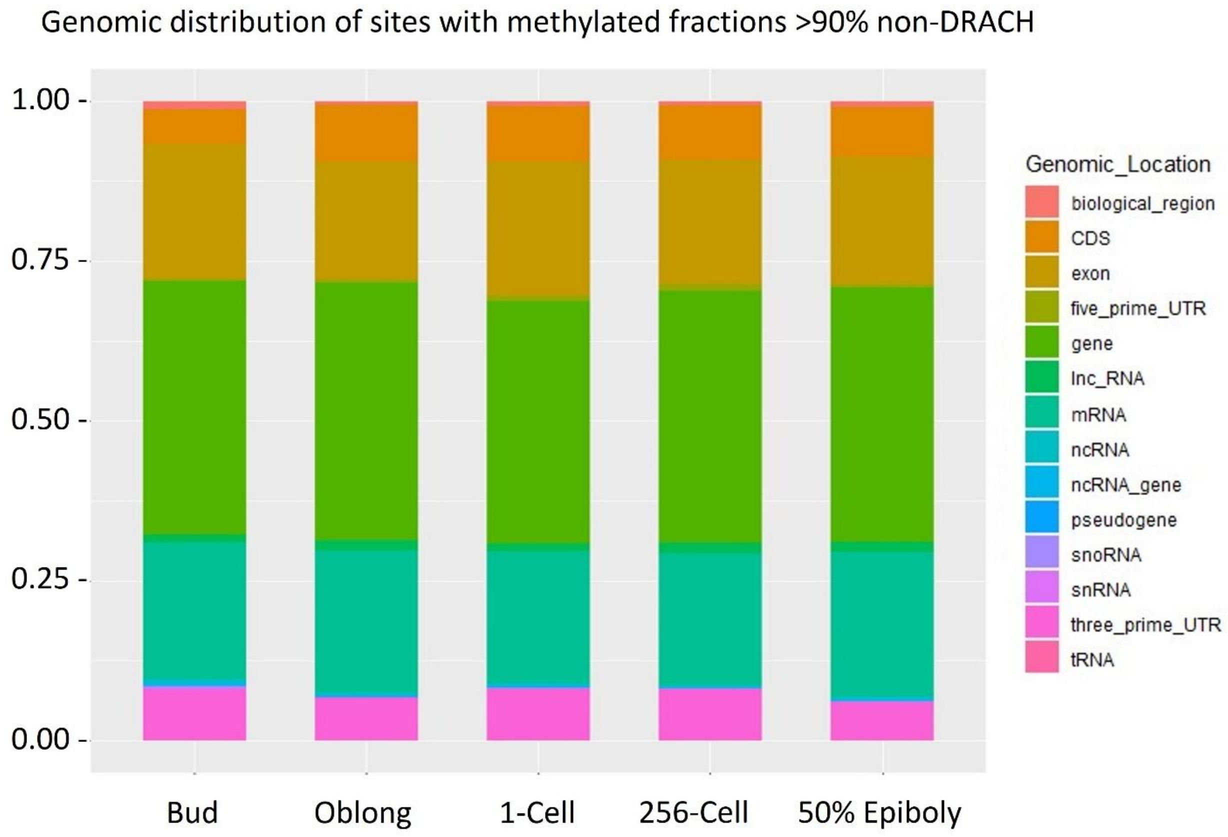


**Supplementary Figure S4:** Genomic distribution of non-DRACH $m^6A$ sites that present 90% - 100% methylated fraction.

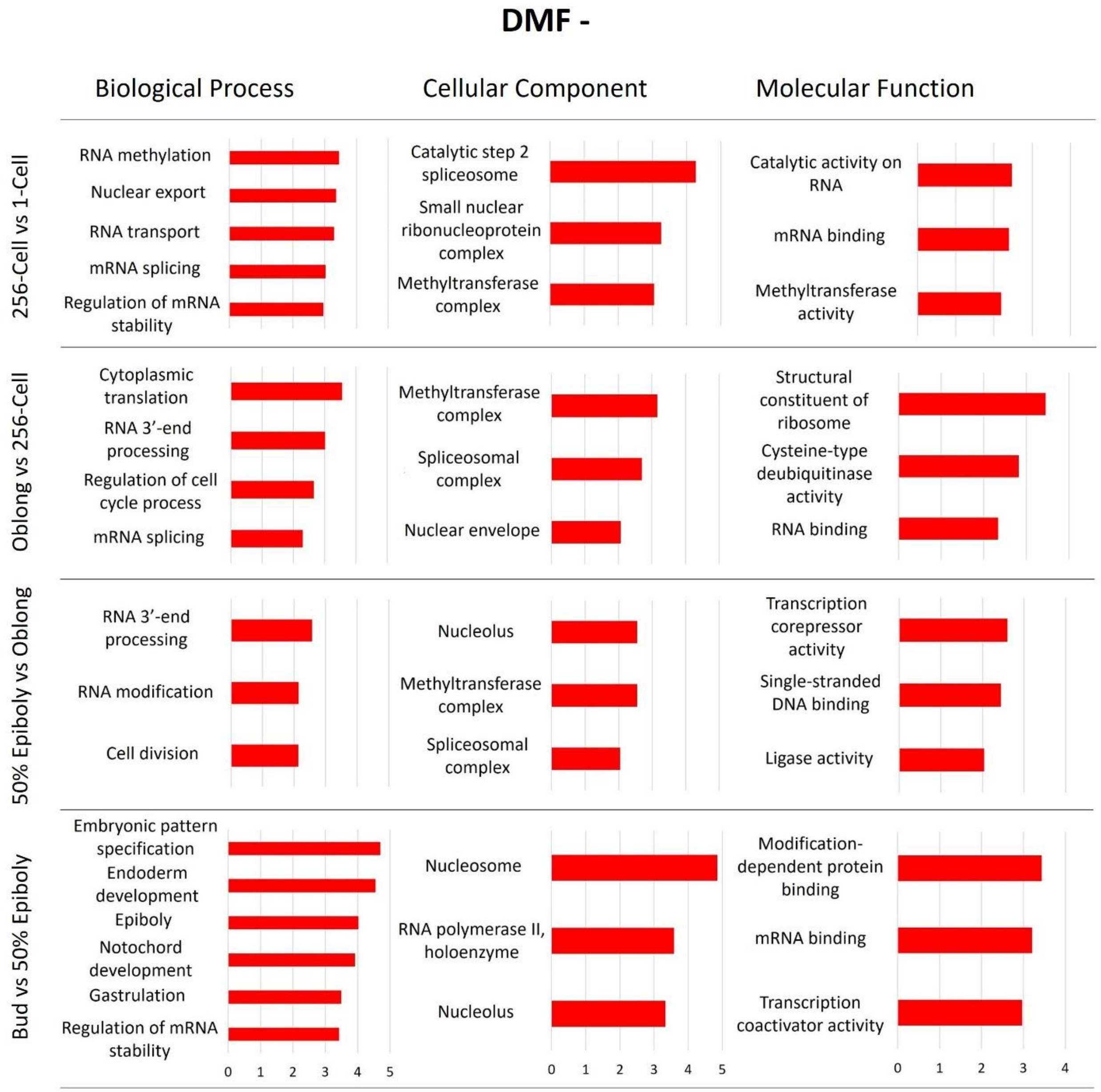


**Supplementary Figure S5:** Gene ontology (GO) analysis results of DMF- genes of the pairwise comparison between zebrafish embryonic stages. DMF refers to genes that presented a differential methylated fraction ≥ 20%, which can be positive (DMF+) or negative (DMF-) differential methylation. Selected categories contain a minimum of 50 reference genes and present a >2-fold enrichment. DMF genes during maternal-to-zygotic transition is considering the differentially methylated fractions calculated by comparing 1-cell and 256-cell to 50% epiboly and bud stages.

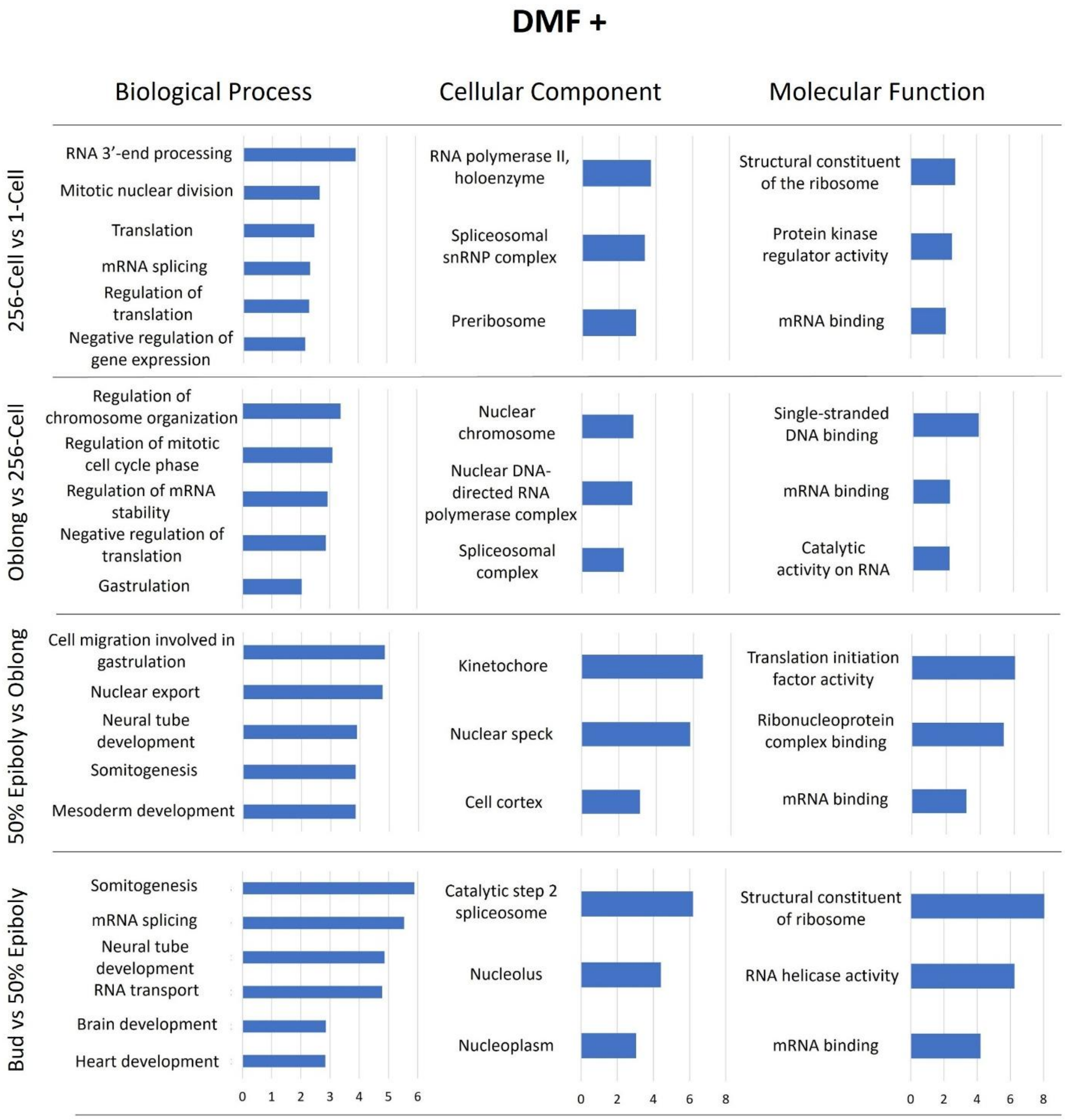


**Supplementary Figure S6:** Gene ontology (GO) analysis results of DMF+ genes of the pairwise comparison between zebrafish embryonic stages.

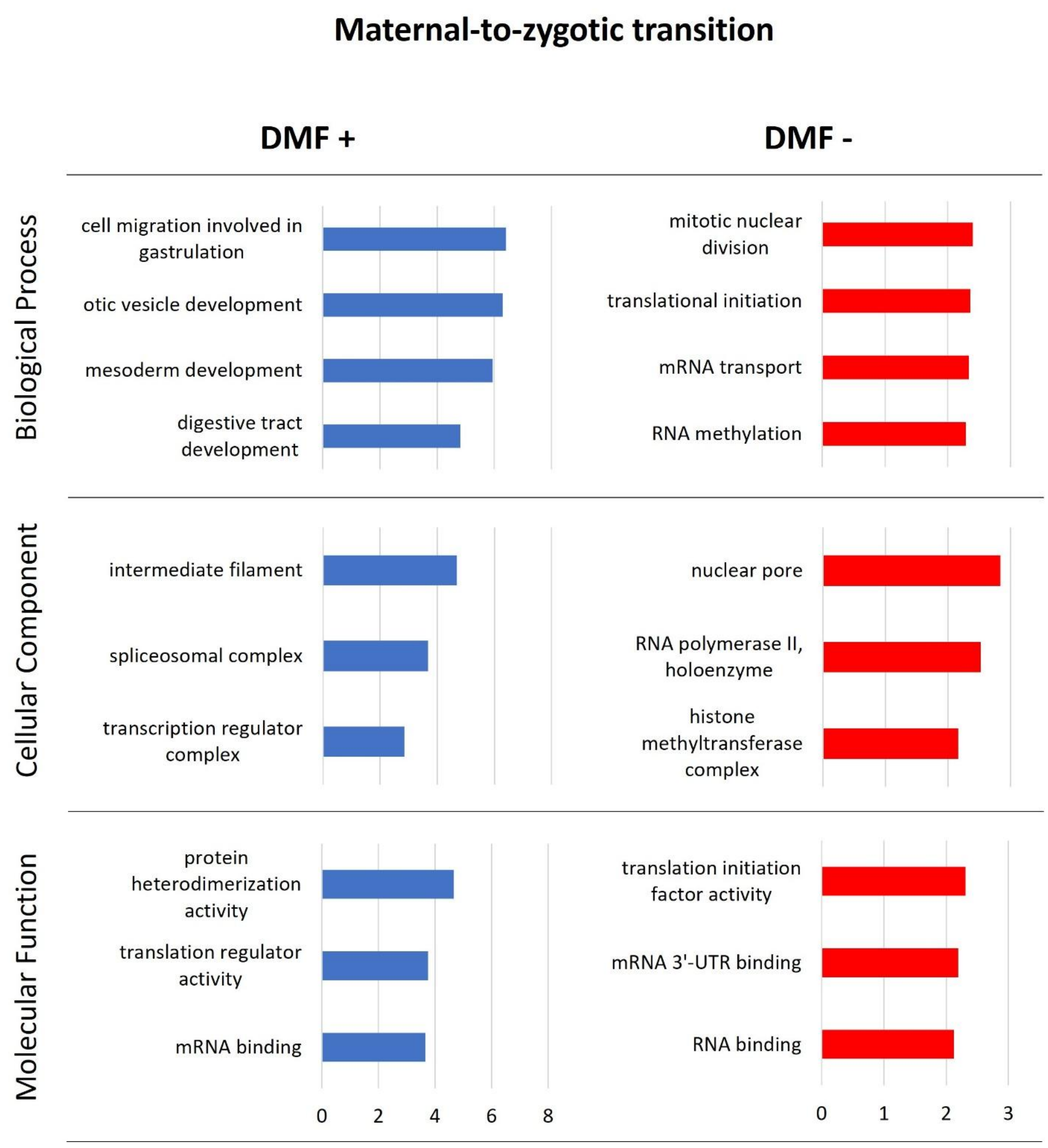


**Supplementary Figure S7:** Gene ontology (GO) analysis results of DMF- and DMF+ genes during maternal-to-zygotic transition.

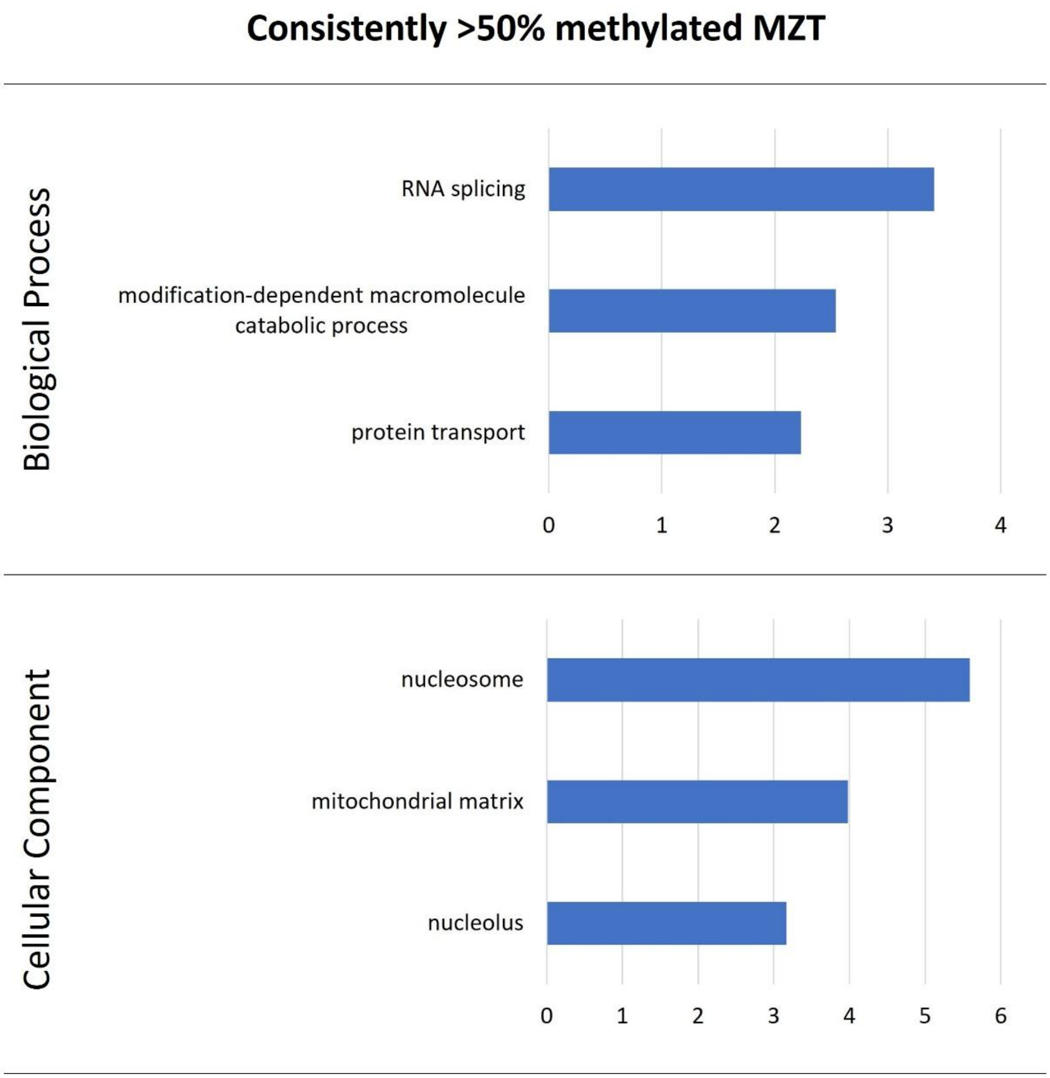


**Supplementary Figure S8:** Gene ontology (GO) analysis results of DMF- and DMF+ genes that were consistently methylated during zebrafish development.

**Supplementary Data File Legends**

**Supplementary Table S1:** RNA concentrations of sampled zebrafish embryos pre and post-ribodepletion.

**Supplementary Table S2:** Number of sequencing reads and mapping values of HiSeq sequencing data for each zebrafish embryonic stage and replicate.

**Supplementary Table S3:** General statistics of $m^6A$ sites mapping to genic regions and mapping uniquely to single transcripts.

**Supplementary File S1:** $m^6A$-SAC-seq data of called $m^6A$ sites for each zebrafish embryonic stage after stringent calling of $m^6A$ sites.

**Supplementary File S2:** List of 147 $m^6A$ locations shared between all 5 developmental stages and its corresponding gene.

**Supplementary File S3:** Global matrix of $m^6A$ methylated fraction per gene per stage in the context of maximum number of sites.

**Supplementary File S4:** Gene ontology results of DMF- genes per paired-stage comparison.

**Supplementary File S5:** Gene ontology results of DMF+ genes per paired-stage comparison.

**Supplementary File S6:** Differential methylation during maternal-to-zygotic transition, calculated by comparing 1-cell and 256-cell to 50% epiboly and bud stages.

**Supplementary File S7:** Gene ontology results of DMF- and DMF+ genes during maternal-to-zygotic transition.

**Supplementary File S8:** Genes that presented a methylated fraction ≥ 50% across zebrafish development.

**Supplementary File S9:** Gene ontology results of a methylated fraction ≥ 50% during zebrafish embryonic development.

**Supplementary File S10:** Gene ontology results of genes that shared identical methylated sites in all five zebrafish embryonic stages.
