## Supplementary Table S1 for "Single-nucleotide resolution mapping of m^6^A of zebrafish mRNAs in early embryonic development links m^6^A modifications to the maternal-to-zygotic transition"

Supplementary Table S1. RNA concentrations of sampled zebrafish embryos pre and post-ribodepletion.

| **Prior to ribodepletion** | | | | Post ribodepletion | |
| --- | --- | --- | --- | --- | --- |
| Sample ID | **Concentration (ng/ul)** | **Total RNA (ug)** | **RINe** | **Ribo- RNA (ng/ul) Tapestation** | **Ribo- RNA (ng/ul) Qubit** |
| 1-cell 1 | 587 | 21.719 | 9.7 | 10.2 | 5.8 |
| 1-cell 2 | 306 | 11.322 | 9.8 | 12.8 | 9.04 |
| 1-cell 3 | 369 | 13.653 | 9.8 | 8.45 | 7.22 |
| 256-cell 1 | 219 | 8.103 | 9.9 | 18 | 10.8 |
| 256-cell 2 | 199 | 7.363 | 10 | 19 | 13.1 |
| 256-cell 3 | 271 | 10.027 | 10 | 16.9 | 12.2 |
| oblong 1 | 227 | 8.399 | 10 | 17.3 | 9.62 |
| oblong 2 | 210 | 7.77 | 9.7 | 16.4 | 11 |
| oblong 3 | 265 | 9.805 | 10 | 20.9 | 13.1 |
| 50% epi 1 | 204 | 7.548 | 10 | 20.3 | 16.3 |
| 50% epi 2 | 212 | 7.844 | 9.9 | 21.7 | 13.8 |
| 50% epi 3 | 205 | 7.585 | 9.7 | 24.6 | 19.2 |
| bud 1 | 210 | 7.77 | 9.9 | 21.1 | 21 |
| bud 2 | 239 | 8.843 | 9.9 | 17.9 | 14.7 |
| bud 3 | 251 | 9.287 | 10 | 16.7 | 9.58 |
