## Supplementary Table S2 for "Single-nucleotide resolution mapping of m^6^A of zebrafish mRNAs in early embryonic development links m^6^A modifications to the maternal-to-zygotic transition"

Supplementary Table S2. Number of sequencing reads and mapping values of HiSeq sequencing data for each zebrafish embryonic stage and replicate.

| **Stage** | Number of reads | Passing cutadapt filter | %mapped reads |
| --- | --- | --- | --- |
| 1-cell_1 | 45.915.184 | 45.190.201 | 56.68 |
| 1-cell_2 | 54.455.391 | 53.879.899 | 54.32 |
| 1-cell_3 | 44.375.861 | 43.891.356 | 42.44 |
| 256-cell_1 | 49.707.024 | 49.219.732 | 50.18 |
| 256-cell_2 | 47.308.102 | 47.069.217 | 39.15 |
| 256-cell_3 | 47.655.521 | 46.608.772 | 48.40 |
| oblong_1 | 47.828.197 | 47.209.907 | 58.55 |
| oblong_2 | 54.193.660 | 53.996.815 | 50.31 |
| oblong_3 | 47.101.503 | 46.759.785 | 27.10 |
| 50%_epiboly_1 | 61.980.117 | 61.521.589 | 42.96 |
| 50%_epiboly_2 | 43.583.665 | 43.108.272 | 41.02 |
| 50%_epiboly_3 | 34.694.479 | 34.296.102 | 42.54 |
| bud_1 | 36.828.955 | 36.510.900 | 40.76 |
| bud_2 | 46.295.690 | 46.092.793 | 44.39 |
| bud_3 | 49.320.234 | 48.650.895 | 30.82 |
| 1-cell_FTO+_1 | 42.769.078 | 42.676.860 | 73.59 |
| 1-cell_FTO+_2 | 36.280.937 | 36.201.819 | 68.14 |
| 1-cell_FTO+_3 | 71.808.598 | 71.527.223 | 56.17 |
| 256-cell_FTO+_1 | 34.751.940 | 34.657.032 | 55.16 |
| 256-cell_FTO+_2 | 113.391.598 | 113.156.974 | 37.10 |
| 256-cell_FTO+_3 | 43.618.476 | 43.451.957 | 61.01 |
| oblong_FTO+_1 | 41.615.414 | 41.446.704 | 61.49 |
| oblong_FTO+_2 | 53.143.320 | 53.005.555 | 62.77 |
| oblong_FTO+_3 | 43.352.290 | 42.870.505 | 52.82 |
| 50%_epiboly_FTO+_1 | 61.329.662 | 60.543.448 | 58.62 |
| 50%_epiboly_FTO+_2 | 61.587.138 | 60.919.612 | 49.27 |
| 50%_epiboly_FTO+_3 | 53.570.606 | 52.837.238 | 52.21 |
| bud_FTO+_1 | 49.608.944 | 49.371.620 | 49.58 |
| bud_FTO+_2 | 50.084.175 | 49.783.085 | 55.06 |
| bud_FTO+_3 | 52.132.262 | 51.867.647 | 49.17 |
