## Supplementary Table S3 for "Single-nucleotide resolution mapping of m^6^A of zebrafish mRNAs in early embryonic development links m^6^A modifications to the maternal-to-zygotic transition"

Supplementary Table S3. General statistics of m⁶A sites mapping to genic regions and mapping uniquely to single transcripts.

| **Stage** | **Total number of m6A sites** | m6A sites mapping to genic regions | m6A sites unique to single transcripts |
| --- | --- | --- | --- |
| 1-cell | 12.104 | 10.479 | 6.483 |
| 256-cell | 15.264 | 12.807 | 7.979 |
| oblong | 13.681 | 11.049 | 6.756 |
| 50% epiboly | 3.531 | 3.039 | 1.798 |
| bud | 2.165 | 1.645 | 1.047 |
